# The developing epicardium regulates cardiac chamber morphogenesis by promoting cardiomyocyte growth

**DOI:** 10.1101/2021.08.09.455639

**Authors:** Giulia L. M. Boezio, Josephine Gollin, Shengnan Zhao, Rashmi Priya, Shivani Mansingh, Stefan Guenther, Nana Fukuda, Felix Gunawan, Didier Y. R. Stainier

## Abstract

The epicardium, the outermost layer of the heart, is an important regulator of cardiac regeneration. However, a detailed understanding of the crosstalk between the epicardium and myocardium during development requires further investigation. Here, we generated three models of epicardial impairment in zebrafish by mutating the transcription factor genes *tcf21* and *wt1a*, and by ablating *tcf21*^+^ epicardial cells. Notably, all three epicardial-impairment models exhibit smaller ventricles. We identified the initial cause of this phenotype as defective cardiomyocyte growth, resulting in reduced cell surface and volume. This failure of cardiomyocytes to grow is followed by decreased proliferation and increased abluminal extrusion. By temporally manipulating its ablation, we show that the epicardium is required to support ventricular growth during early cardiac morphogenesis. By transcriptomic profiling of sorted epicardial cells, we identified reduced expression of FGF and VEGF ligand genes in *tcf21*^-/-^ hearts, and pharmacological inhibition of these signaling pathways partially recapitulated the ventricular growth defects. Thus, the analysis of these epicardial-impairment models further elucidates the distinct roles of the epicardium during cardiac morphogenesis and the signaling pathways underlying epicardial-myocardial crosstalk.

## Introduction

The epicardium is the last layer to incorporate into the heart during development. Epicardial cells (EpiCs) delaminate from the extra-cardiac proepicardial organ (PEO) and attach to the naked myocardium as free-floating cells due to the physical properties of the pericardial fluid (Rodgers et al., 2008; Peralta et al., 2013). The epicardium forms a mesothelial layer that completely envelops the heart, then undergoes epithelial-to-mesenchymal transition (EMT) and gives rise to various epicardial derived cells (EPDCs) (Smith et al., 2011; Acharya et al., 2012; Smits et al., 2018). While the epicardium becomes dormant after undergoing EMT, it reactivates after cardiac injury and upregulates developmental genes as well as new gene regulatory networks (Weinberger et al., 2021), and it rapidly regenerates (Masters and Riley, 2014; Cao and Poss, 2018).

The epicardium has received great attention due to its ability to differentiate into a multitude of cell types during cardiac repair and to its role as a source of paracrine signals that promote wound healing (Masters and Riley, 2014; Cao and Poss, 2018). However, a detailed understanding of the epicardial-myocardial crosstalk during development has proven more elusive. As the few identified factors involved in epicardial-myocardial signaling, including components of the fibroblast growth factor (FGF) and insulin growth factor (IGF) signaling pathways, are expressed in developmental and regenerative contexts (Pennisi et al., 2003; Lavine et al., 2005; Brade et al., 2011; Li et al., 2011; Vega-Hernandez et al., 2011), identifying the processes underlying epicardial-myocardial crosstalk during development has important implications for cardiac repair.

Defects in epicardial coverage consistently result in a common myocardial phenotype—small, underdeveloped ventricles. Ablation of the PEO in chicken embryos causes reduced cardiac size and occasional ventricular bulging (Manner, 1993; Pennisi et al., 2003; Manner et al., 2005; Takahashi et al., 2014). Similarly, mutations in several epicardial-enriched genes, including those encoding the transcription factors TCF21 and WT1, abrogate epicardial coverage, leading to a reduction in ventricular size (Moore et al., 1999; Acharya et al., 2012). Most studies to date have concluded that the major role of the epicardium is to promote CM proliferation (Pennisi et al., 2003; Lavine et al., 2005; Li et al., 2011) and to contribute to the ventricular mass by giving rise to EPDCs such as fibroblasts (Mahtab et al., 2009; Acharya et al., 2012). Notably, a few studies have started to challenge the view that the sole function of the epicardium is to regulate CM cell cycle (Eid et al., 1992; Kastner et al., 1994; Takahashi et al., 2014), but they have so far been limited to using *in vitro* explants or fixed tissue sections. Deeper investigation of epicardial function in promoting myocardial growth requires a model in which the cellular phenotypes can be experimentally followed in 4 dimensions.

Here, we generated three models of epicardial impairment in zebrafish larvae by mutating the transcription factor genes *tcf21* and *wt1a*, and by ablating EpiCs. Leveraging the advantages of these newly established models and the amenability of zebrafish to live imaging at 3D resolution, we identified a novel role for the epicardium in promoting CM growth and determined the time-window when this epicardium to myocardium interaction occurs. We also generated a transcriptomic dataset of epicardial-enriched factors to identify molecules important for this crosstalk. Focusing on *fgf24* and *vegfaa*, we provide evidence that they are epicardial-enriched regulators of ventricular growth.

## Results

### Generation of three zebrafish epicardial-impairment models

In zebrafish, EpiCs start adhering to the myocardial wall around 52-56 hours post fertilization (hpf), and cover the entire ventricle by 96-120 hpf (Fig. 1A) (Peralta et al., 2014). To study the epicardial-myocardial crosstalk, we generated three different zebrafish models with impaired epicardial coverage. First, we mutated *tcf21* and *wt1a* (Fig. S1C), two transcription factor genes enriched in the epicardium (Fig. S1A, B) (Serluca, 2008; Liu and Stainier, 2010; Peralta et al., 2014)). Our *tcf21* deletion allele contains a frameshift in the coding sequence, leading to a predicted truncated protein with an incomplete DNA-binding domain (Fig. S1C), while the mutation does not affect the stability of the mutant mRNA (Fig. S1 D). Conversely, our *wt1a* promoter deletion leads to a complete absence of *wt1a* mRNA (Fig. S1C, E). In mouse, the lack of either transcription factor leads to impaired epicardial coverage, but its impact on myocardial development is poorly understood (Moore et al., 1999; Acharya et al., 2012). We observed that in zebrafish, mutations in *tcf21* and *wt1a* also lead to a reduction of *TgBAC(tcf21:*NLS-EGFP)^+^(hereafter referred to as *tcf1*^+^) EpiCs on the ventricular wall, as evidenced at 54 hpf and -even more prominently- at 76 and 100 hpf (Fig. 1B-H). However, whereas *wt1a* mutants exhibit a complete absence of *tcf21^+^* EpiCs on the ventricular wall, the epicardial coverage reduction in *tcf21* mutants is variable (Fig. 1I). This phenotypic variability is likely not due to the *tcf21* mutation leading to a hypomorphic allele, as non-cardiac phenotypes previously identified in *tcf21* mutants, including the lack of head muscles (Nagelberg et al., 2015; Burg et al., 2016), are observed with complete penetrance in our *tcf21* mutants (n>300 larvae). Notably, the number of outflow tract (OFT) *tcf21^+^* EpiCs appears unaffected in both mutants (Fig. 1J), likely due to the different origin of this epicardial population (Perez-Pomares et al., 2003; Weinberger et al., 2020). Second, to establish a model in which epicardial coverage can be impaired in a specific time window, we used the previously described nitroreductase/metronidazole (NTR/MTZ) system (Curado et al., 2007; Pisharath et al., 2007; Curado et al., 2008). By treating *TgBAC(tcf21:mCherry-NTR*) (Wang et al., 2015) embryos (NTR^+^) with MTZ, we could ablate nearly all *tcf21*^+^ cells before the epicardium covers the ventricle (52-100 hpf; Fig. 1K-M), thereby establishing an inducible system which complements our two mutant models. Using the pan-epicardial marker Caveolin1 (Cao et al., 2016), we confirmed that *TgBAC(tcf21:* NLS-EGFP)expression is a reliable marker for all EpiCs, and that the loss of *tcf21^+^* cells in our models is due to an absence of EpiCs and not to the loss of *TgBAC(tcf21:*NLS-EGFP) expression. Caveolin1 immunostaining was only present in “escaper” ventricular *tcf21*^+^ EpiCs in *tcf21*^−^ hearts, and around the OFT in all three models (Fig. S1F-I’). Altogether, these three distinct epicardial-impairment models constitute a complementary set of reagents to study the effects of epicardial impairment on cardiac morphogenesis. Moreover, using the genetic models generated here as well as others, it will be interesting to investigate how Wt1a and Tcf21 regulate epicardial development.

**Figure 1:**
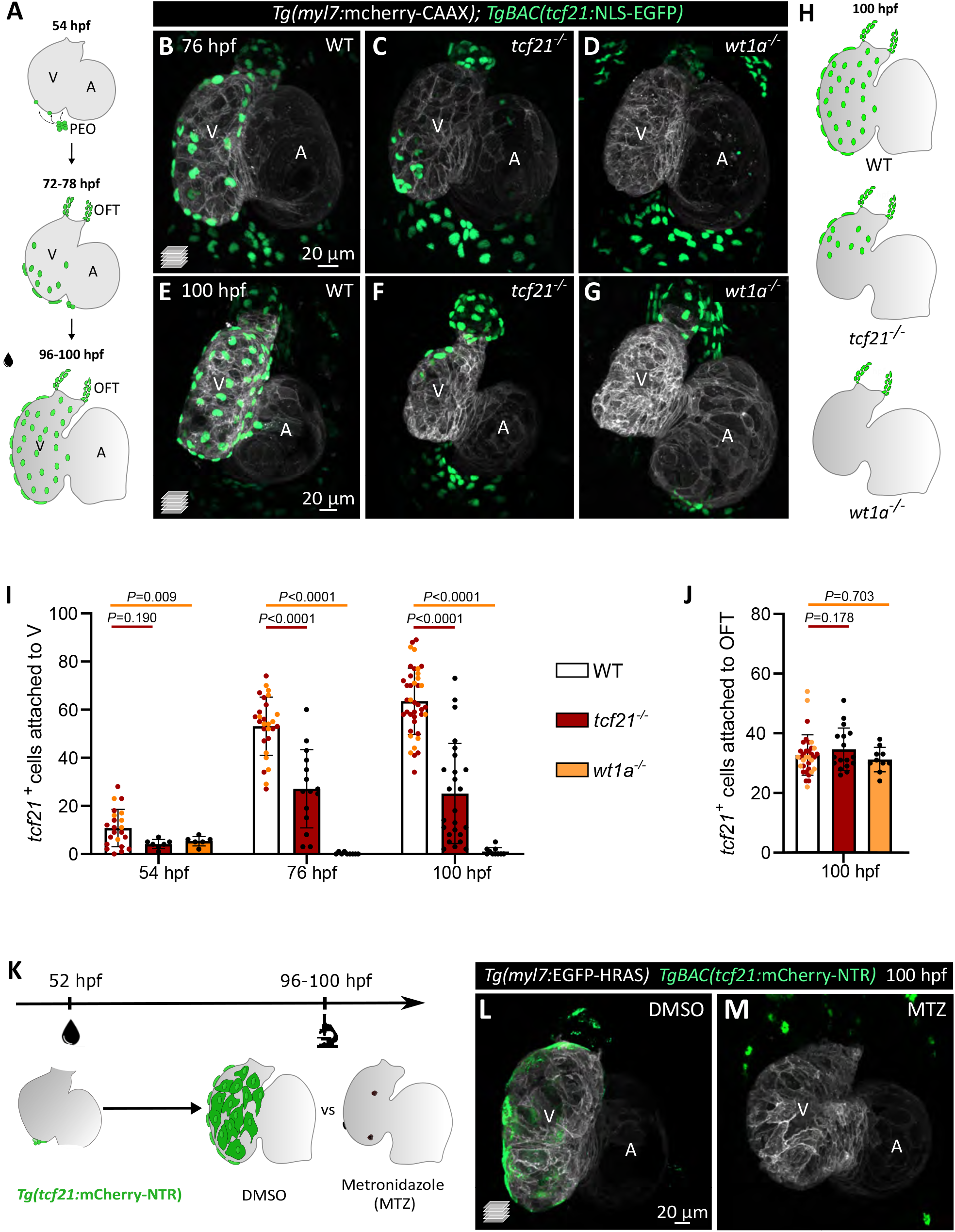
The transcription factors tcf21 and Wt1a are required for epicardial attachment to the ventricle. **A**) Schematic representation of the epicardial coverage of the zebrafish embryonic and larval heart. **B-G**) Confocal images of 76 (B-D) and 100 (E-G) hpf *TgBAC(tcf21:NLS-EGFP); Tg(myl7:mCherry-CAAX)* larvae. **H**) Schematics of the epicardial coverage in 100 hpf WT, *tcf21*^-/-^ and *wt1a*^-/-^ larvae. Grey, myocardium; green, EpiCs. **I-J**) Quantification of *tcf21*^+^ EpiCs attached to the ventricular myocardium (H) and the OFT (I). The colors of WT dots refer to *tcf21*^+/+^ (red) and *wt1a*^+/+^ (orange) siblings. Mean ± SD; *P* values from *t*- or Mann-Whitney tests (following normality test) compared with +/+ siblings of each genotype. **K**) Epicardial ablation protocol using the NTR/MTZ system. **L-M**) Confocal images of 100 hpf *TgBAC(tcf21:mCherry-NTR); Tg(myl7:EGFP-HRAS)* hearts, showing the absence of EpiCs post MTZ treatment (M), compared with DMSO-treated larvae (L). WT, wild types; A, atrium; V, ventricle; OFT, outflow tract; PEO, proepicardial organ.

### Impairment in cardiomyocyte growth becomes evident before reduced cardiomyocyte proliferation when epicardial cells are lost

We next aimed to determine how epicardial impairment affects ventricular morphogenesis. Starting at 96 hpf, we observed pericardial edema in *tcf21*^-/-^, *wt1a*^-/-^ and NTR^+^ MTZ-treated larvae (Fig. S1J-M), together with impaired ventricular fractional shortening starting at 100 hpf (Fig. S2A). The cardiac ventricle in all epicardial-impairment models was also approximately 30% smaller than in wild type (WT; Fig. 2A-D, S2B-E). Interestingly, from 76 to 100 hpf, we observed that the wild-type ventricle grew on average by 36.4% in volume, whereas the mutant ventricle started with a comparable volume at 76 hpf but failed to enlarge over time (Fig. 2E).

**Figure 2:**
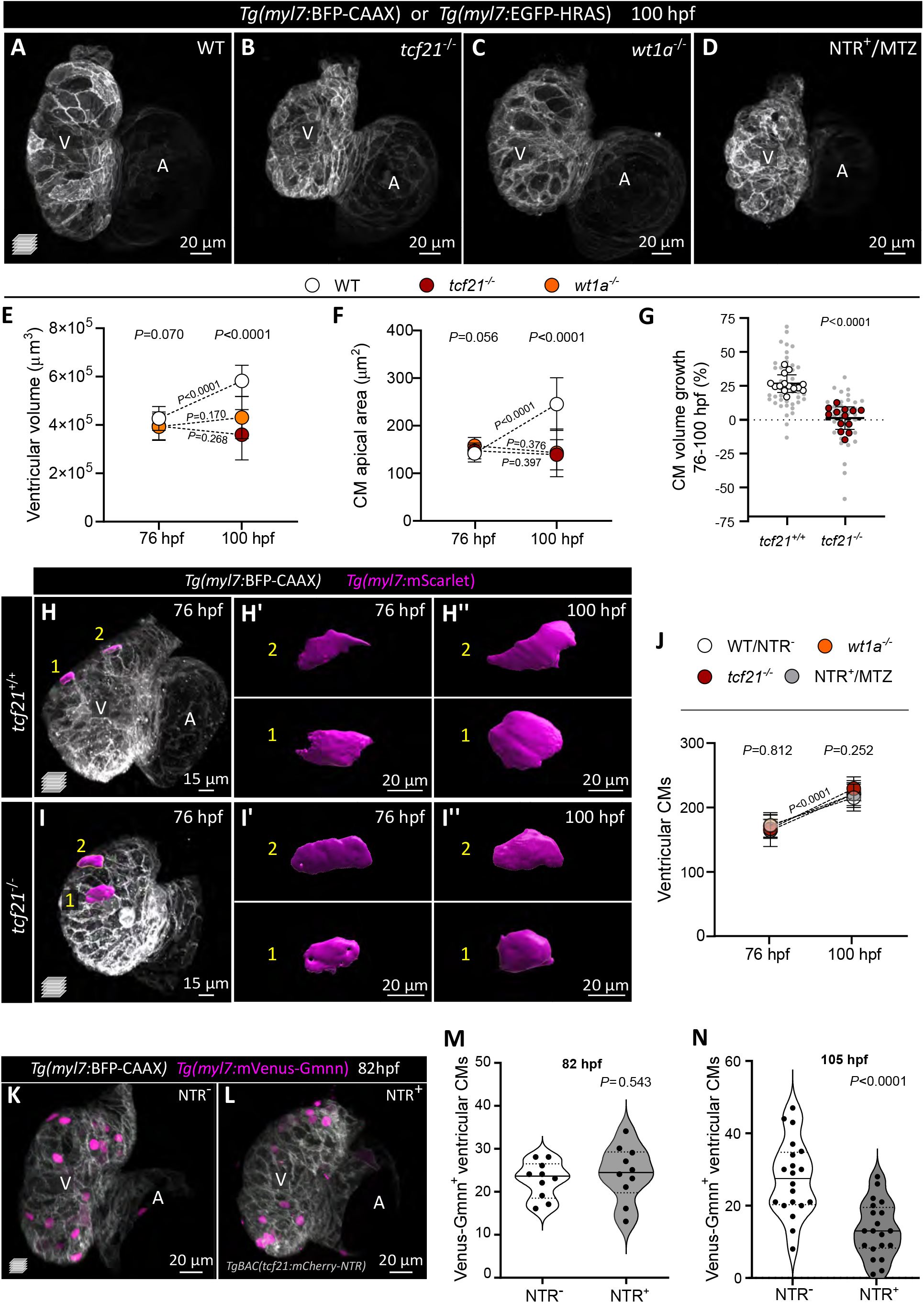
Impaired epicardial coverage affects ventricular cardiomyocyte size increase and ventricular growth. **A-D)** Confocal images of 100 hpf WT, *tcf21*^-/-^, *wt1a*^-/-^, and *TgBAC(tcf21:mCherry-NTR)*^+^(NTR^+^) MTZ-treated larvae, exhibiting reduced ventricular size. **E, F**) Change in ventricular volume (E) and CM apical area (F) from 76 to 100 hpf in WT, *tcf21*^-/-^ and *wt1a*^-/-^ larvae. **G**) Measurement of individual CM volume increase in percentage between 76 and 100 hpf, as measured through the *myl7*:mScarlet signal; large dots, average per larva; small dots, individual CMs. Mean ± SD; *P* values from *t*-test comparing the averages per larva. **H-I”**) Confocal images of *Tg(myl7:BFP-CAAX); tcf21*^+/+^ or *tcf21*^-/-^ larva at 76 and 100 hpf (same larvae shown), injected with *myl7:mScarlet* DNA to label individual CMs; (H’-I’’) 3D surface rendering of individual CMs at the two time points. **J**) Changes in ventricular CM numbers from 76 to 100 hpf in WT (or NTR^-^), *tcf21*^-/-^, *wt1a*^-/-^ and NTR^+^-MTZ treated larvae. **K-N**) Confocal images and quantification of *myl7:*mVenus-Gmnn^+^ CMs in 82 (K-M) and 105 (N) hpf control (NTR^-^) and NTR^+^ MTZ-treated larvae. **E, F, J**) Mean ± SD; *P* values from One-Way ANOVA amongst the three different genotypes at the same time-point (above the graph), or *t*-tests comparing the two different time-points within the same genotype (on the dotted lines). Single data points are shown in Figure S2. **M, N**) Median (solid line) and quartiles (dashed lines); *P* values from *t*-tests. WT, wild types; A, atrium; V, ventricle.

Increase in organ size is driven by hypertrophic (increase in cell size) and hyperplastic (increase in cell number) growth. The first phenotype we observed between control and epicardial-impairment models was in the CM apical area. Although the average apical area of compact layer CMs was comparable in WT and mutant larvae at 76 hpf, it was significantly smaller in mutant CMs compared to WT at 100 hpf (Fig. 2F, S2F). To assess the volumetric growth of individual compact layer CMs over time, we tracked single CMs in *tcf21*^+/+^ and *tcf21*^-/-^ hearts by mosaic expression of *Tg(myl7:mScarlet)*. Strikingly, *tcf21*^+/+^ CM volume increased by 26.8% between 76 and 100 hpf, while *tcf21*^-/-^ CM volume did not significantly change (+1.4%) (Fig. 2G-I). Our data are, to our knowledge, the first to correlate an increase in CM cell volume with ventricular growth, and to uncover a requirement for the epicardium in promoting CM cell growth during cardiac development.

While the epicardium has not been previously linked with CM hypertrophic growth, it has been implicated in promoting CM proliferation (Pennisi et al., 2003; Lavine et al., 2005; Li et al., 2011). We therefore assessed the number of proliferating CMs at 82 hpf, a time point before the growth defects can be observed, by counting the number of Venus-Gmnn^+^ CMs (cells in the S/G/M phases (Sugiyama et al., 2009; Choi et al., 2013). We observed no significant difference between control and NTR^+^ MTZ-treated larvae (Fig. 2K-M). We also counted the number of ventricular CMs, and they increased by a similar proportion (≈30%) in WT larvae and in larvae from all three models between 76 and 100 hpf (Fig. 2J), resulting in comparable numbers of CMs at all time points analyzed (Fig. S2G). Interestingly, at 105 hpf, a time point subsequent to the appearance of CM size defects, we observed a severe reduction in the number of Venus-Gmnn^+^ CMs in NTR^+^ MTZ-treated larvae compared with controls (−48.5%; Fig. 2N). These defects in CM proliferation may thus be a consequence of the initial impairment in CM growth. In support of this interpretation, eukaryotic cell cycle progression is known to depend on cell growth (Jorgensen and Tyers, 2004), and a multitude of cell types, including CMs, expand in size prior to dividing (Son et al., 2015; Zlotek-Zlotkiewicz et al., 2015; Uribe et al., 2018). In addition, we observed CM extrusion away from the cardiac lumen in the three epicardial-impairment models (Fig. S3A-F’), consistent with previous reports (Manner et al., 2005; Rasouli et al., 2018). We hypothesize that in the absence of the epicardium, excessive cellular density -as observed by reduced inter-nuclear distance in the epicardial-impairment models (Fig. S3G-I)-drives the aberrant extrusion of a few CMs. However, this CM extrusion in the epicardial-impairment models only plays a minor role in their reduced ventricular size.

Altogether, our observations uncover a previously unidentified role for the epicardium in promoting the initial stages of CM growth, which subsequently affects CM proliferation and ventricular growth.

### Epicardial cells are required and sufficient for ventricular cardiomyocyte growth in a restricted early time window

To further analyze the dependency of CM growth on the epicardium, we tested whether rescuing the epicardial coverage was sufficient to improve ventricular growth. Leveraging the temporal versatility of the NTR/MTZ system, we ablated the epicardium specifically from 52 to 100 hpf and then washed out the MTZ; we first confirmed that the EpiCs recover by 144 hpf (Fig. 3A-C), as previously reported (Wang et al., 2015). Strikingly, epicardial restoration was sufficient to ameliorate the cardiac growth defects in MTZ-treated larvae (Fig. 3B’-D).

**Figure 3:**
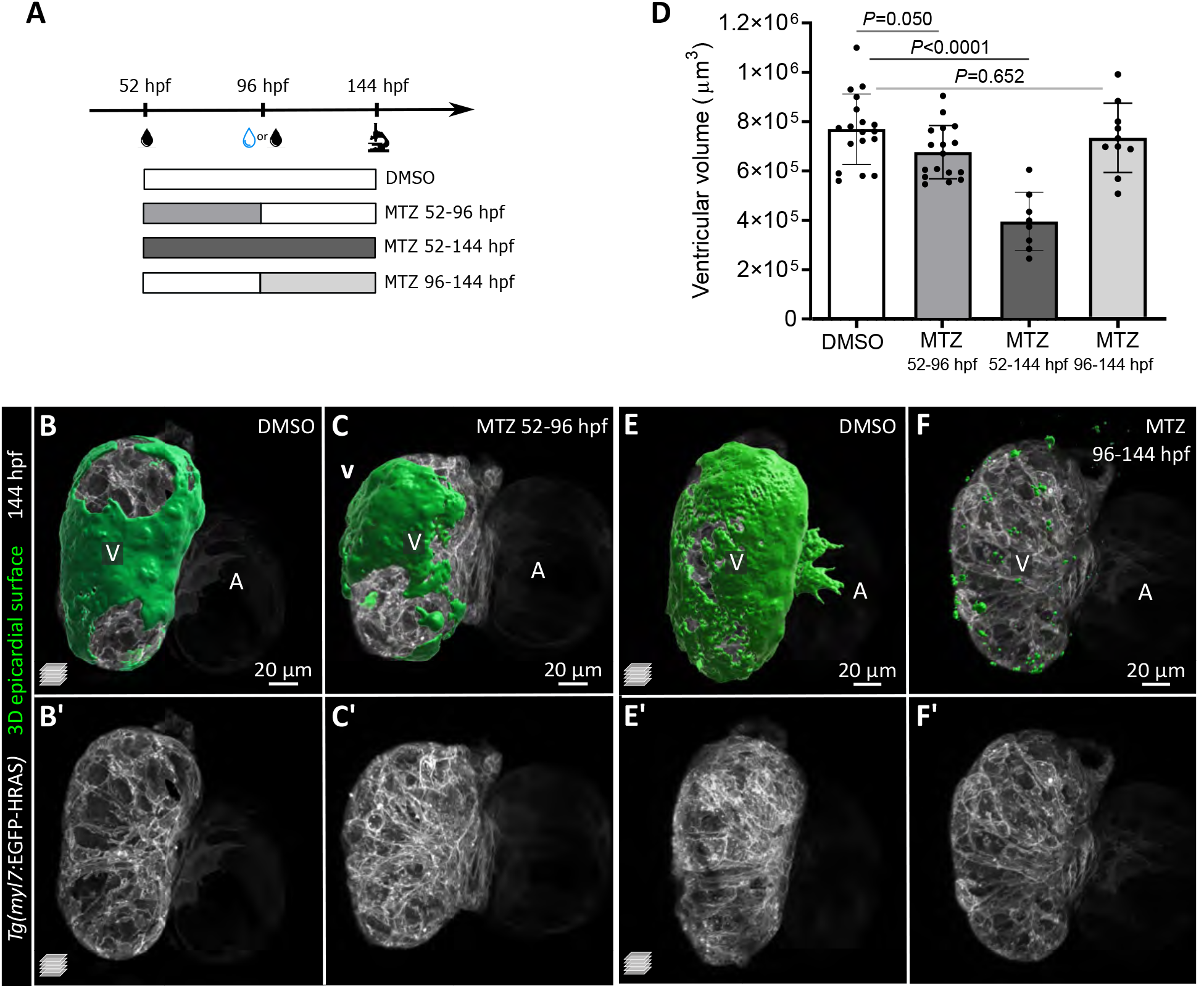
Epicardial cells are required for ventricular cardiomyocyte growth during early cardiac morphogenesis, but are dispensable at later timepoints. **A**) Schematic of the MTZ treatment protocol. **B-C’**) Confocal images of 144 hpf *Tg(myl7: EGFP-HRAS); TgBAC(tcf21:mCherry-NTR)* DMSO and MTZ-treated larvae. Green, 3D reconstruction of epicardial coverage, showing a partial recovery of EpiCs in 52-96 hpf MTZ treated larvae. **D**) Quantification of ventricular volume at 144 hpf following epicardial regeneration or late epicardial ablation. **E-F’**) Confocal images of 144 hpf *Tg(myl7:EGFP-HRAS); TgBAC(tcf21:mCherry-NTR)* DMSO and MTZ-treated larvae. Green, 3D reconstruction of epicardial coverage, showing the lack of epicardium in 96-144 hpf MTZ-treated larvae. **D**) Mean ± SD; *P* values from One-Way ANOVA test. A, atrium; V, ventricle.

Previous studies investigating the consequences of reduced epicardial coverage have used genetic models or physical ablation of the PEO, but the constitutive lack of epicardium fails to pinpoint the time window in which EpiCs promote myocardial development. As mentioned above, we identified the 52-100 hpf window to be crucial for epicardial-myocardial interactions, which coincides with the period of epicardial attachment. We then tested the effects of epicardial ablation between 96 and 144 hpf (Fig. 3A); surprisingly, while epicardial ablation appeared quite effective (Fig. 3 E, F), we observed no obvious morphological defects in ventricular morphology or size (Fig. 3D-F’).

These results suggest that epicardial-myocardial crosstalk is necessary to regulate ventricular volume during 52-100 hpf, but dispensable once epicardial coverage is complete (96 hpf). We propose that, from 96 hpf onwards, the CMs continue to grow due to epicardial-independent intrinsic or extrinsic cues. Therefore, at later stages, the epicardium might assume a different function, including preparing for the onset of EMT. Future investigations of later epicardial function during cardiac development (e.g., at the onset of EMT and EPDCs formation) will greatly benefit from the temporal versatility of this NTR/MTZ model.

### Epicardial-derived secreted factors promote ventricular cardiomyocyte growth

Next, we aimed to understand how the epicardium modulates CM growth. The epicardium is an important signaling center during development and regeneration (Quijada et al., 2020). Nonetheless, the appearance of extruding CMs in epicardial-deficient larvae (Fig. S3), which was also observed following PEO ablation in chick (Manner et al., 2005), raised questions as to whether the epicardium primarily acts as a physical barrier to preserve myocardial integrity. To further investigate the role of the epicardium as a mechanical support and/or a signaling source, we focused on the sizable fraction of *tcf21*^-/-^ hearts that exhibited substantial epicardial coverage on their ventricle (Fig. 1H; Fig. S4A-C). We first observed that these mutants still displayed defects in ventricular size that were comparable with those in *tcf21*^-/-^ hearts devoid of epicardial coverage (Fig. S4A-C). We found no significant correlation between the number of ventricular EpiCs and ventricular volume (Fig. S4D), or the number of extruding CMs (Fig. S4F). Notably, we also observed that some extruding CMs were covered by EpiCs (Fig. S4, E’), suggesting that the physical presence of EpiCs alone does not prevent CM extrusion.

To identify epicardial factors necessary for CM growth, we compared the transcriptomes of sorted *tcf21*^+/+^ and *tcf21*^-/-^ EpiCs and CMs at 96 hpf (Fig. 4A; Supplementary Tables 2 and 3). To minimize bias in the analyses, we selected *tcf21*^-/-^ larvae that exhibited a wild-type-like ventricular epicardial coverage and collected the same number of EpiCs from the two genotypes. We first analyzed the genes expressed in the two wild-type populations by RNA-seq and confirmed the expression of cell-specific markers, including *postnb, wt1a, col1a2, cav1, aldh1a2* in EpiCs and *ttn.1, ttn.2, myh7, myh6* in CMs.

**Figure 4:**
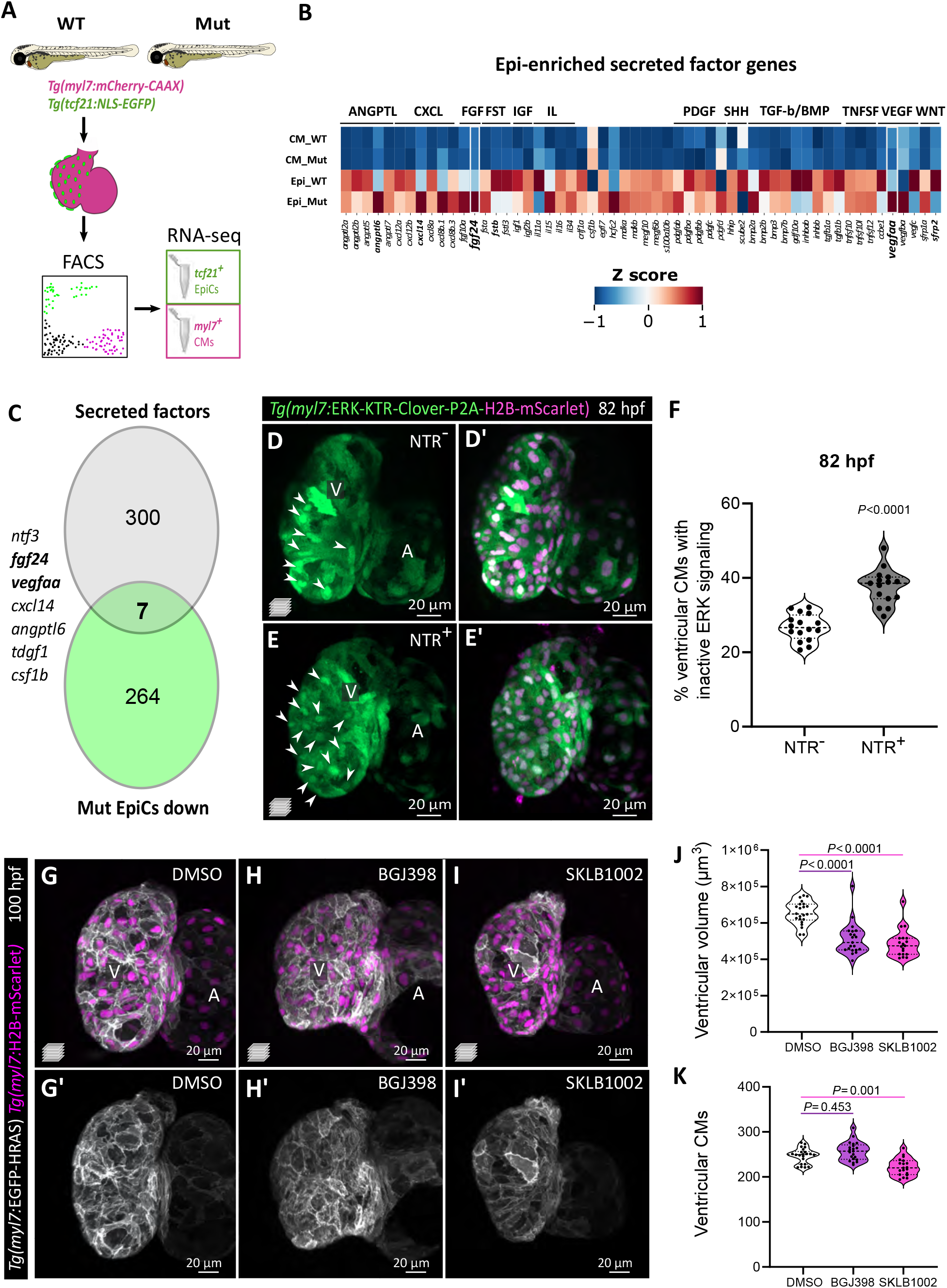
Epicardial-enriched *FGF* and *VEGF* ligand genes in epicardial-myocardial crosstalk. **A**) RNA-seq from *tcf21*^+^ and *myl7*^+^ cells from 96 hpf *tcf21*^+/+^ and *tcf21*^-/-^ hearts. **B**) Heatmap of zebrafish secreted factor genes in *tcf21*^+/+^ and *tcf21*^-/-^ EpiCs and CMs, showing Z scores normalized per row. The genes highlighted in bold are differentially expressed between *tcf21*^+/+^ and *tcf21*^-/-^ EpiCs (log2FC < |0.7|, *P* adj.<0.05). **C**) Venn diagram denoting the intersection between secreted factor genes in the zebrafish genome and genes that are downregulated (log2FC < −0.7, *P* adj.<0.05) in *tcf21*^-/-^ EpiCs. **D-F**) Confocal images (D, E) and quantification (F) of 82 hpf *Tg(myl7:ERK-KTR-Clover-P2A-H2B-mScarlet)* ventricles in control (NTR^-^) and *TgBAC(tcf21:mCherry-NTR)* (NTR^+^) larvae treated with MTZ. White arrowheads point to CMs with nuclear Clover (inactive ERK), quantified in F. **G-K**) Confocal images (G-I’), quantification of ventricular volume (J) and CM numbers (K) of 100 hpf larvae treated with FGFR (BGJ398) and VEGFR (SKLB1002) inhibitors starting at 65 hpf. A, atrium; V, ventricle.

We focused on epicardial-derived secreted factors that potentially mediate this epicardial-myocardial crosstalk. Amongst the factors enriched in the EpiCs compared to CMs are members of the FGF, IGF, transforming growth factor (TGF)-β, and platelet-derived growth factor (PDGF) pathways (Fig. 4B; Supplementary Table 3). These pathways are important for epicardial-myocardial crosstalk in mammals (Olivey and Svensson, 2010; Li et al., 2017), suggesting that the molecular regulators of this crosstalk are similar in zebrafish.

Amongst the downregulated secreted factors in *tcf21*^-/-^ EpiCs are *FGF* and *VEGF* ligand genes, including *fgf10a, fgf24, vegfaa, vegfba* (Fig. 4B, C; Supplementary Table 3). *fgf24* and *vegfaa* (Fig. 4C, S5A), in particular, are the *FGF* and *VEGF* ligand genes with the highest epicardial expression in our dataset and the only ones significantly downregulated (*P* adj. <0.05; Supplementary Table 3). The FGF pathway mediates epicardial-myocardial crosstalk in mouse and chicken embryos, where it is primarily known for its role in promoting CM proliferation (Pennisi et al., 2003; Lavine et al., 2005). By *in situ* hybridization, we observed *fgf24* expression in 76 hpf hearts including in the epicardium (Fig. S5B, B’). On the other hand, Vegfaa promotes angiogenesis and coronary vessel formation (Liang et al., 2001; Wu et al., 2012; Marin-Juez et al., 2016; Rossi et al., 2016), but its role in the developing epicardium has not been investigated until recently (Bruton et al., 2021). We observed that *vegfaa* expression in the developing heart appears to be initially limited to the epicardium until 100 hpf (Fig. S5C-D”; (Bruton et al., 2021)), at which point it also becomes expressed in the endocardium (Karra et al., 2018). The epicardial enrichment of *fgf24* and *vegfaa* (log2FC: +4,77 and +3,38, respectively), as well as their downregulation in *tcf21*^-/-^ EpiCs led us to hypothesize that epicardial-derived Fgf24 and Vegfaa mediate signaling to the myocardium. To test this hypothesis, we first assessed the activation of the mitogen-activated protein kinase (MAPK)/extracellular regulated kinase (ERK) pathway, a downstream effector of both FGF and VEGF signaling, in CMs. To do so, we generated a transgenic line, *Tg(myl7:ERK-KTR-Clover-P2A-H2B-mScarlet)*, to monitor ERK activation through a kinase translocation reporter (de la Cova et al., 2017; Mayr et al., 2018; Okuda et al., 2020) (Fig. 4D). Following MTZ treatment between 52 and 80 hpf, *TgBAC(tcf21:mCherry-NTR);Tg(myl7:ERK-KTR-Clover-P2A-H2B-mScarlet)* hearts devoid of epicardium exhibited a significantly increased number of CMs with inactive ERK signaling compared with control hearts (Fig. 4D-F). In addition, we used BGJ398 (De Simone et al., 2021) and SKLB1002 (Zhang et al., 2011) to inhibit the FGF and VEGF signaling pathways, respectively, from 65 to 100 hpf. Larvae treated with these compounds recapitulated the smaller ventricle phenotype. FGF inhibition affected ventricular volume without causing any changes in CM number, whereas the VEGF inhibitor led to a mild but significant decrease in ventricular CMs (−10%; Fig. 4G-K). It is likely that a global inhibition of the VEGF pathway leads to a stronger phenotype compared to the epicardial-specific downregulation of *vegfaa* in *tcf21*^-/-^ hearts, and/or that SKLB1002 affects additional signaling pathways.

Interestingly, the mammalian ortholog of Fgf24 binds Fgfr4 (Mok et al., 2014), which is highly expressed in zebrafish CMs and downregulated in *tcf21*^-/-^ CMs, as per our transcriptomic datasets (Supplementary Table 2). We speculate that the downregulation of Fgfr4 in CMs might be caused by feedback loops caused by the reduction in Fgf24. On the other hand, Vegfaa is not known to bind receptors prominently expressed in CMs, but binds Vegfr2/Kdrl, which is enriched in EpiCs. Therefore, Vegfaa potentially signals in an autocrine manner, similar to retinoic acid (Stuckmann et al., 2003; Brade et al., 2011), and regulates the production of other signaling molecules. Otherwise, it was recently proposed that the epicardial expression of *vegfaa* (in response to macrophage activation) regulates Notch activity in the endocardium, which -in turn-signals to CMs (Bruton et al., 2021). Further studies will address the molecular mechanisms by which these signaling pathways mediate epicardial-myocardial crosstalk to promote ventricular morphogenesis.

Alternatively, we cannot exclude the possibility that ECM-related components secreted by the epicardium play a role in maintaining CM homeostasis and promoting their growth. In particular, we observed the downregulation of several ECM component genes, including *col6a1/2, col4a1/2, lama5* and *hapln1a/b*.

Altogether, our data uncover a previously unappreciated requirement for the epicardium in promoting CM growth at the cellular and tissue levels, which takes place prior to its previously reported role in stimulating CM proliferation. Moreover, we provide evidence that this inter-tissue crosstalk is mediated by the FGF and VEGF pathways. We propose that the three epicardial-impairment models used in this study provide genetically tractable, and in one case temporally manipulable, systems that complement existing models to deepen our understanding of the cellular and molecular processes involved in epicardial-myocardial crosstalk during cardiac development.

## Data and Software Availability

The RNA-seq data reported in this paper have been deposited in the Gene Expression Omnibus (GEO) database https://www.ncbi.nlm.nih.gov/geo/query/acc.cgi?acc=GSE174505 (accession GSE174505). (Reviewer token: wbydmsugdfmfhch).

## Acknowledgments

This work was supported by funds from the Max Planck Society to D.Y.R.S., a European Molecular Biology Organization (EMBO) Advanced Fellowship (ALTF 642-2018) and a Canadian Institute for Health Research Fellowship (293898) to F.G., and an EMBO fellowship (LTF 1569-2016), a Humboldt fellowship and a Cardio-Pulmonary Institute Grant (EXC 2026, project ID 390649896) to R.P. We would like to thank Matteo Perino for help with the RNA-seq analyses and critical comments on the manuscript, Ann Atzberger and Khrievono Kikhi for help with the FACS experiments, Michelle Collins, Alessandra Gentile, Srinivas Allanki and Hadil El-Sammak for valuable discussions and comments on the manuscript, Dr. Radhan Ramadass for expert help with microscopy, Helen Allmendinger for experimental assistance, and all the fish facility staff for technical support.

## Author contributions

Conceptualization, G.L.M.B, F.G., and D.Y.R.S.; Methodology, G.L.M.B, F.G., N.F., S.G., R.P., and S.M; Validation, G.L.M.B.; Formal Analysis, G.L.M.B. and S.G.; Investigation, G.L.M.B., S.Z., J.G., and S.G.; Writing – Original Draft, G.L.M.B., F.G., and D.Y.R.S.; Writing – Reviewing & Editing, all; Visualization, G.L.M.B.; Supervision, F.G. and D.Y.R.S; Project Administration, D.Y.R.S.; Funding Acquisition, D.Y.R.S.

---

**Figure S1:**
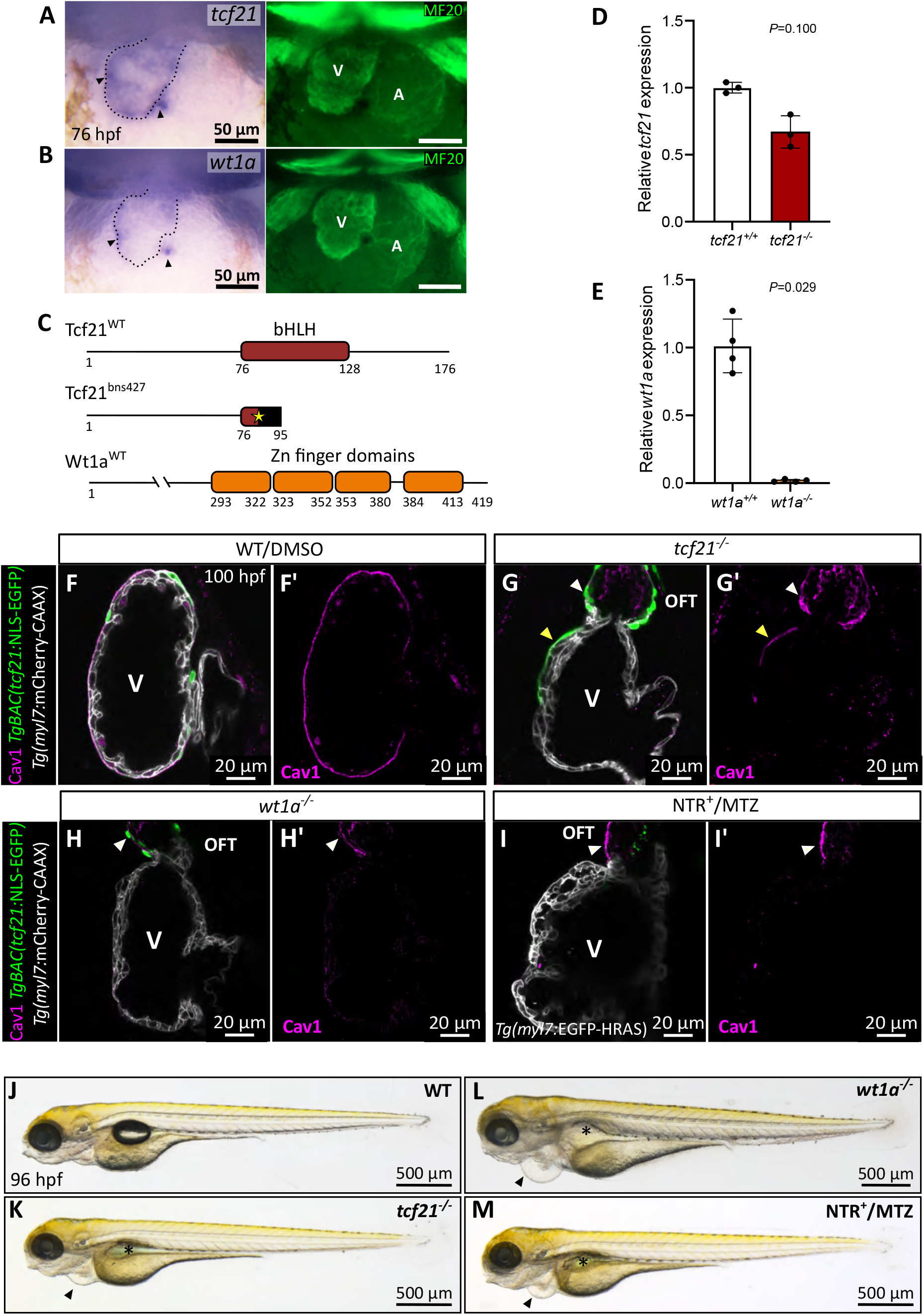
Transcription factors Tcf21 and Wt1a are crucial for cardiac development. **A**, **B**) *In situ* hybridization showing the expression of *tcf21*(*A*) and *wt1a* (B) in 76 hpf hearts. MF20 immunostaining (green) labels the myocardium (dashed lines); arrowheads point to the presumptive signal in the epicardium. **C**) Schematics of wild-type Tcf21 and Wt1a proteins and predicted Tcf21 mutant protein, highlighting the bHLH (red, Tcf21) and Zn finger (orange, Wt1a) domains. Yellow star indicates the CRISPR/Cas9-induced mutation site; the black rectangle represents new sequence downstream of the frameshift-inducing mutation. The *wt1a* mutation targets the promoter region and does not affect the coding sequence. **D, E**) *tcf21* (D) and *wt1a* (E) mRNA levels in 96 hpf *tcf21*^+/+^, *tcf21*^-/-^, *wt1a*^+/+^, and *wt1a*^-/-^ larvae; means ± SD; *P* values from Mann Whitney tests; Ct values are listed in Supplementary Table 1. **F-I’**) Confocal images of 100 hpf hearts immunostained for Caveolin1 (Cav1). Cav1 immunostaining is present only in “escaper” ventricular *tcf21*^+^ EpiCs (yellow arrowheads, *tcf21*^-/-^) and in the OFT (white arrowheads). **J-M**) 96 hpf *tcf21*^-/-^, *wt1a*^-/-^, and NTR^+^ MTZ-treated larvae exhibit pericardial edema (arrowheads) and lack of swim bladder inflation (asterisks). A, atrium; V, ventricle; OFT, outflow tract; bHLH, basic helix–loop–helix; Zn, zinc.

**Figure S2:**
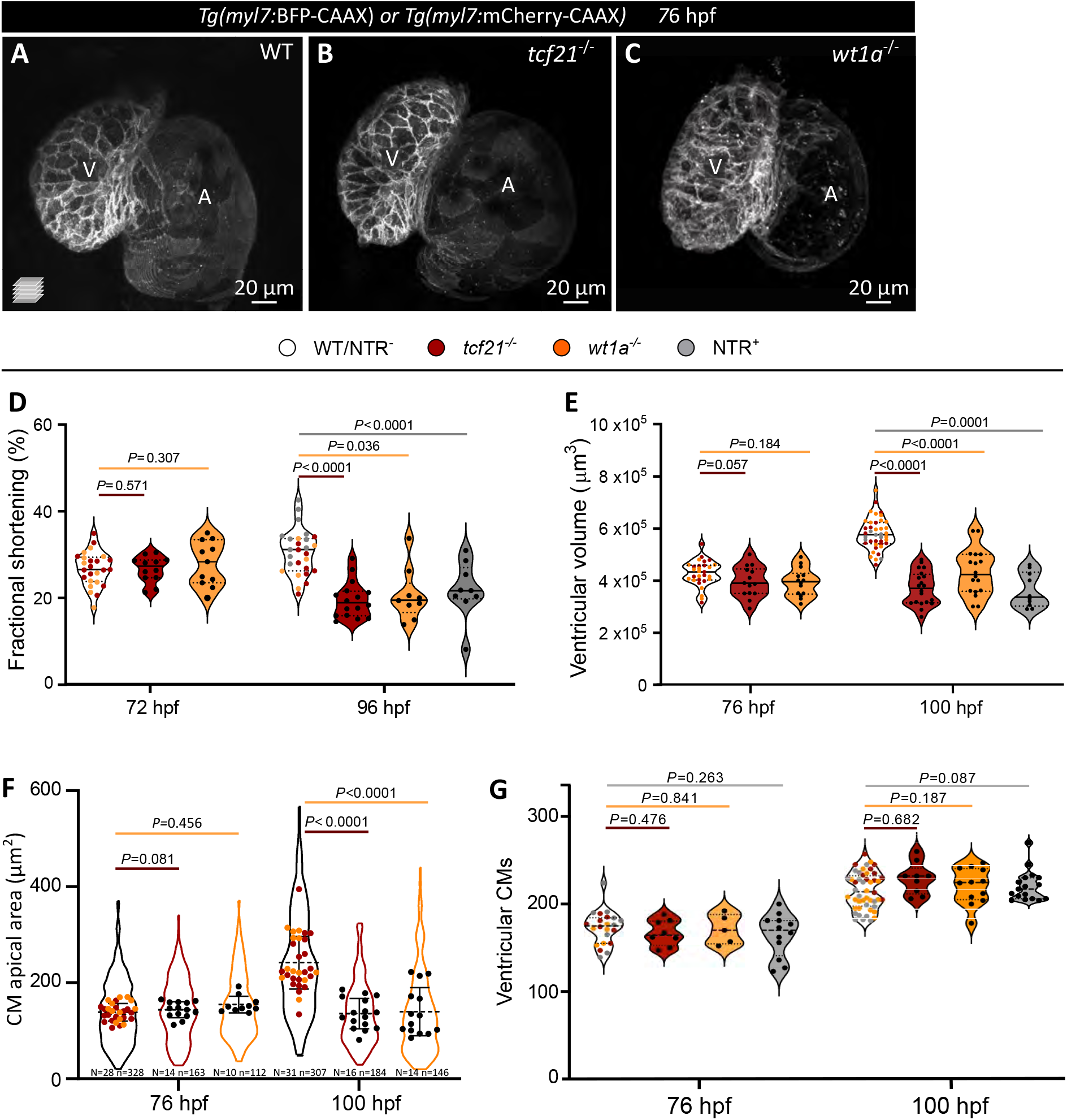
Epicardial impairment affects ventricular size, but not ventricular cardiomyocyte number. **A-C**) Confocal images of 76 hpf WT, *tcf21*^-/-^, and *wt1a*^-/-^ larvae, all exhibiting a similar ventricular size. **D-E**) Quantification of fractional shortening (D) and ventricular volume (E) in WT, mutant and NTR^+^ MTZ-treated larvae; the graph in E relates to Figure 2E, showing the single data points. **F**) CM apical area in WT, *tcf21*^-/-^, and *wt1a*^-/-^ larvae; related to Figure 2F. Violin plot represents the distribution of individual CMs (n); dots represent the average per larva (N). **G**) Ventricular CM numbers in WT, mutant and NTR^+^ MTZ-treated larvae, showing individual data points; related to Figure 2J. **A, B, F, G**) The colors of wild-type dots refer to *tcf21*^+/+^ (red), *wt1a*^+/+^ (orange), and NTR^+^ MTZ-treated (grey) siblings. Median and quartiles (A, B, G), or mean ± SD (F); *P* values from *t*- or Mann-Whitney tests (following normality test), compared with the WT/control siblings of each genotype/treatment. WT, wild types; A, atrium; V, ventricle; N, number of larvae; n, number of CMs.

**Figure S3:**
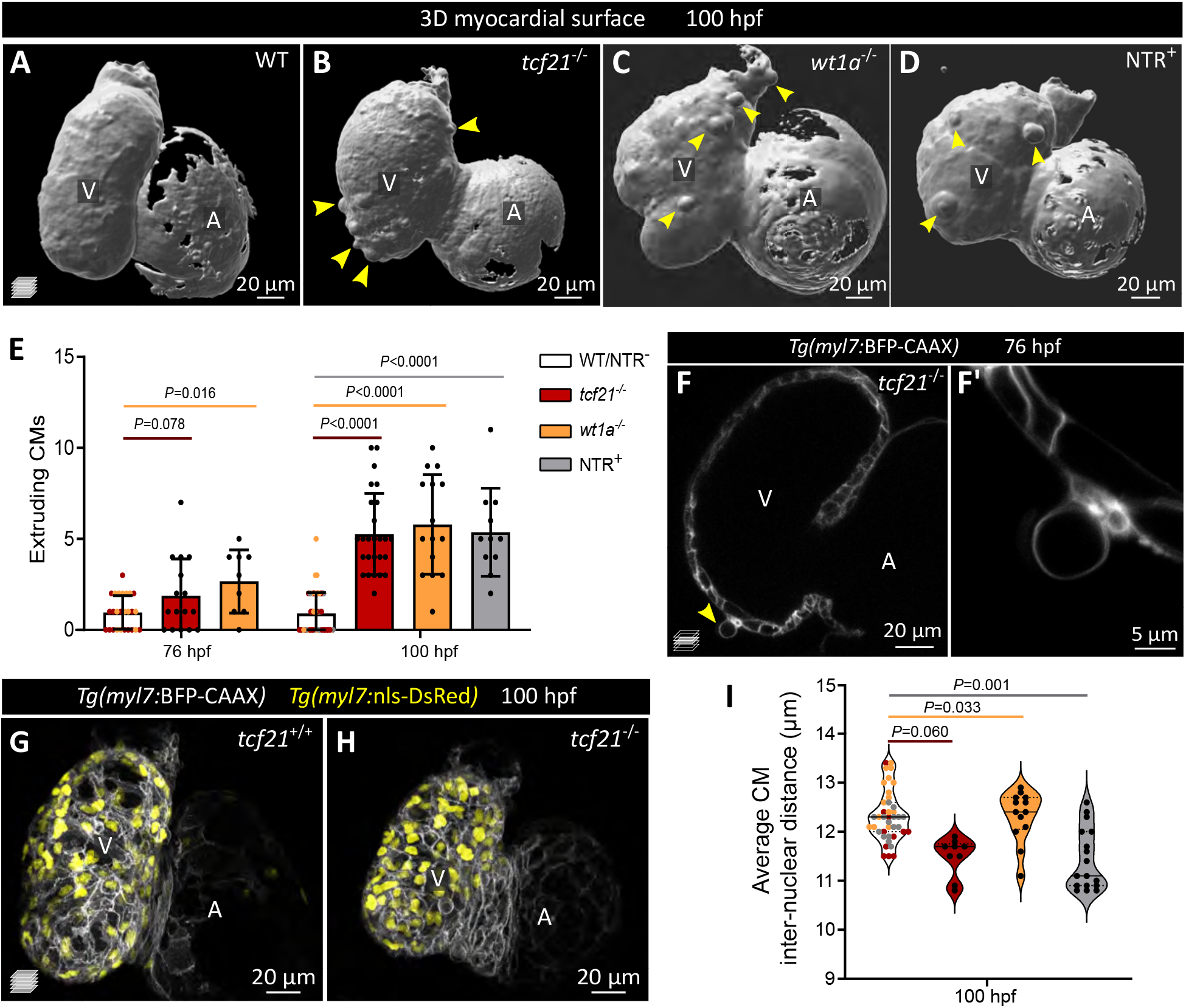
Impaired epicardial coverage causes abluminal cardiomyocyte extrusion. **A-D**) 3D surface rendering of 96 hpf *Tg(myl7:mCherry-CAAX)* (A-C) and *Tg(myl7:EGFP-HRAS)* (D) hearts. Arrowheads point to extruding CMs. **E**) Quantification of CM extrusions at 76 and 96 hpf. **F, F’**) Single-plane images of 76 hpf *Tg(myl7:mCherry-CAAX) tcf21*^-/-^ heart. Arrowheads point to extruding CMs. **G-I**) Confocal images and quantification of the CM inter-nuclear distance in 100 hpf *Tg(myl7:BFP-CAAX);(myl7:nlsDsRed)* hearts. **E, I**) The colors of wild-type dots refer to *tcf21*^+/+^ (red) or *wt1a*^+/+^ (orange) siblings, or NTR^+^ MTZ-treated (grey) siblings. **E, I**) Mean ±SD (E) or median and quartiles (I); *P* values from *t*- or Mann-Whitney tests (following normality test), compared with WT/control siblings of each genotype/treatment. A, atrium; V, ventricle.

**Figure S4:**
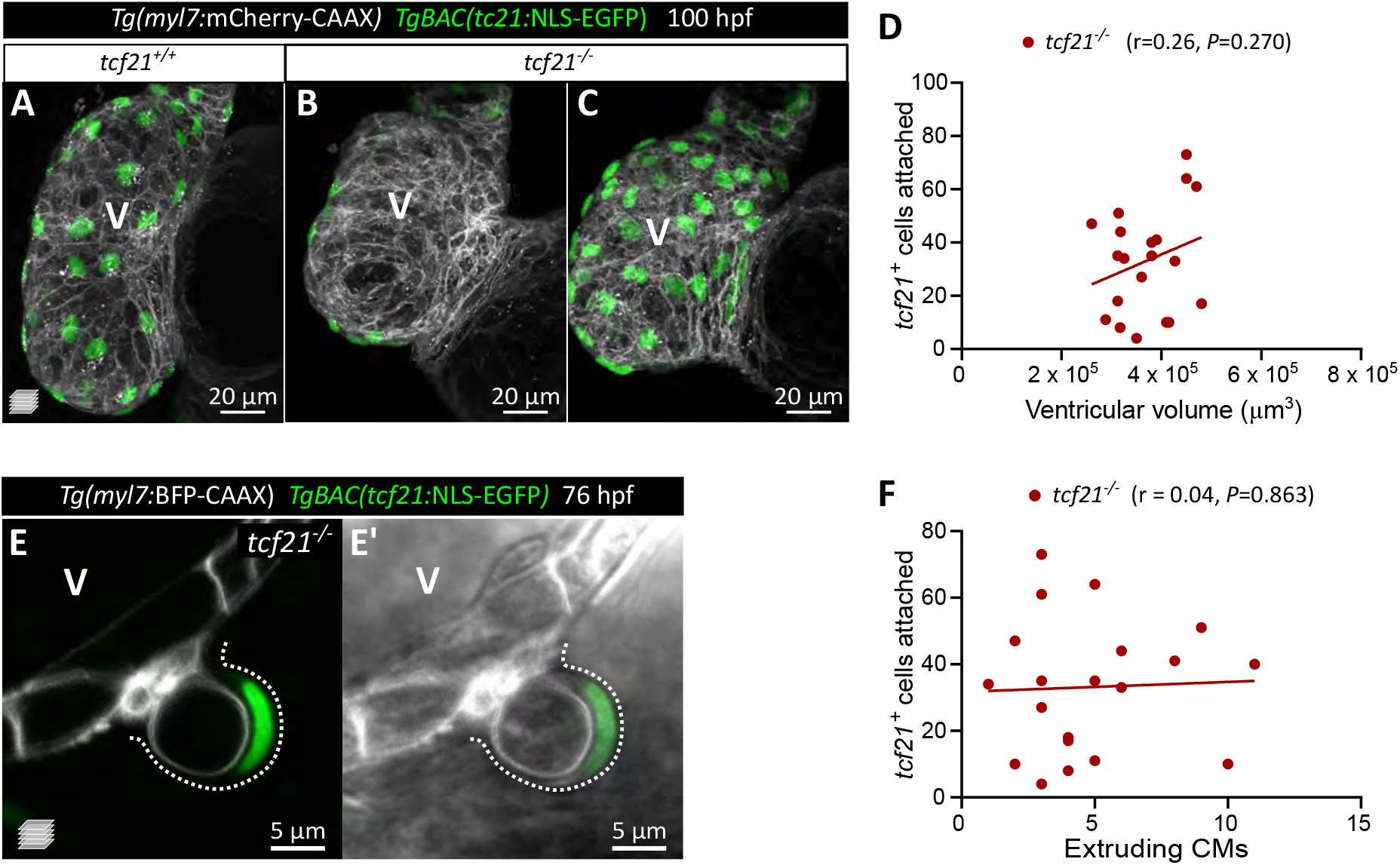
An intact epicardium is required to promote ventricular growth and prevent cardiomyocyte extrusion. **A-C**) Confocal images of 96 hpf *TgBAC(tcf21:NLS-EGFP); Tg(myl7:BFP-CAAX) tcf21*^+/+^ (A) and *tcf21*^-/-^ (B, C) hearts, with different degrees of epicardial coverage (green). **D, F**) Pearson correlation between ventricular volume (D) and extruding CMs (F) (X axis), and the number of ventricular *tcf21*^+^ cells (Y axis) in 96 *tcf21*^-/-^ larvae. **E, E’**) Single confocal plane of a 76 hpf *TgBAC(tcf21:NLS-EGFP); Tg(myl7:mCherry-CAAX) tcf21*^-/-^ ventricle, exhibiting an extruding CM covered by a *tcf21*^+^ epicardial cell (nucleus, green; cell body highlighted with dashed line).

**Figure S5:**
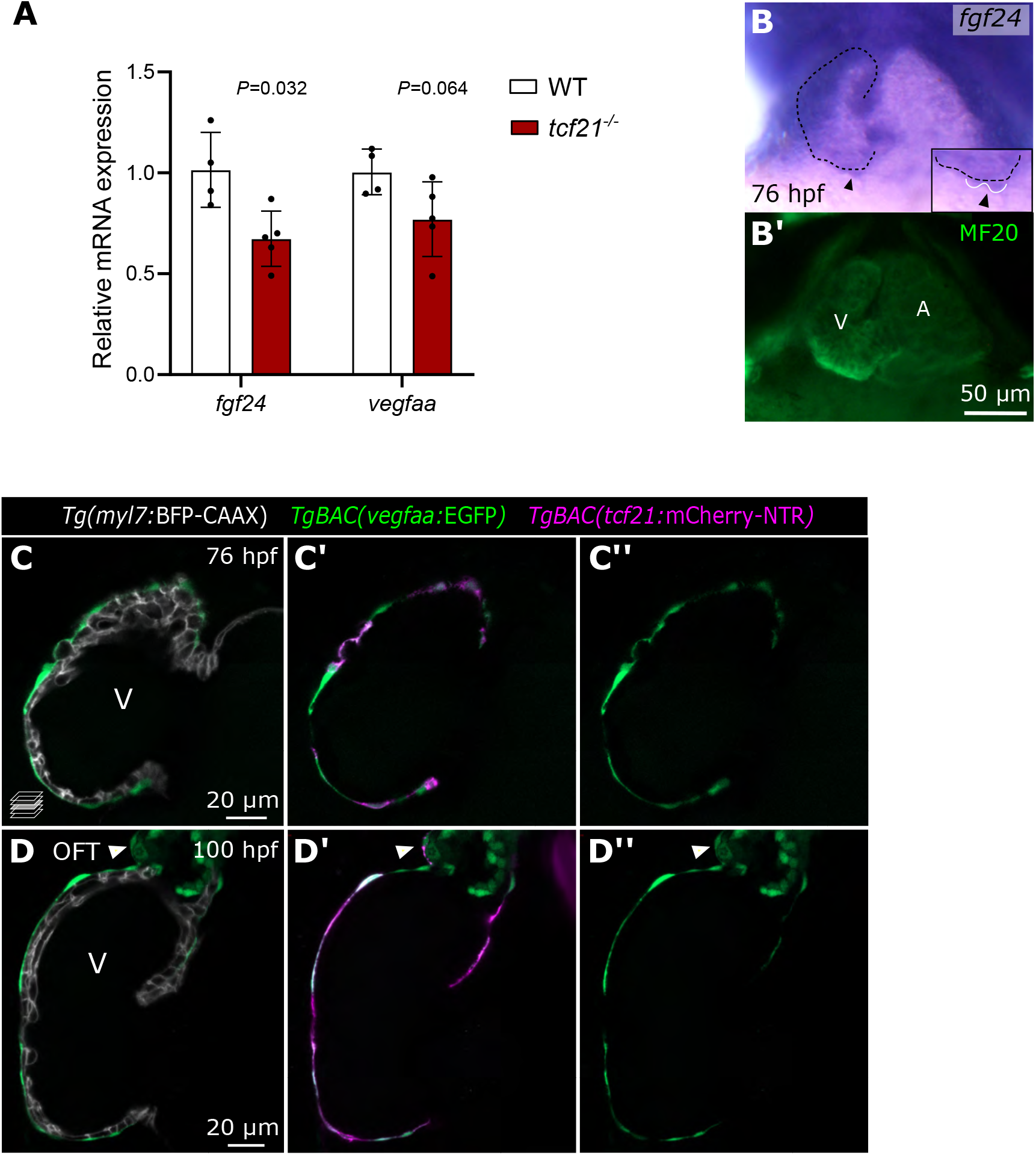
*fgf24* and *vegfaa* expression is enriched in epicardial cells and downregulated in *tcf21*^-/-^ hearts. **A**) *fgf24* and *vegfaa* mRNA levels obtained by RT-qPCR on extracted 96 hpf *tcf21*^+/+^ and *tcf21*^-/-^ hearts; means ± SD; *P* values from Mann Whitney tests; Ct values are listed in Supplementary Table 1. **B, B’**) *In situ* hybridization showing *fgf24* expression in 76 hpf hearts (n=11/11). MF20 immunostaining (green) labels the myocardium (dashed lines). Arrowheads, epicardial cells outside of the myocardial wall (enlarged in the box). **C-D’’**) Confocal images of 76 (C) and 96 (D) hpf *TgBAC(vegfaa:EGFP); Tg(myl7:BFP-CAAX); TgBAC(tcf21:mCherry-NTR)* ventricles. *TgBAC(vegfaa:EGFP)* expression is restricted to the epicardium (here seen on the ventricle and OFT) as well as OFT SMCs (white arrowheads). A, atrium; V, ventricle; OFT, outflow tract.

**Supplementary Table 1: Ct values of genes by RT-qPCR and primers.**

**Supplementary Table 2: List of the top differentially expressed genes (>1 or <−1 log2FC, baseMean > 50) from RNA-seq dataset of 96 hpf CMs sorted from *tcf21*^+/+^ and *tcf21*^-/-^ larval hearts.**

**Supplementary Table 3: List of the top differentially expressed genes (>1 or <−1 log2FC, baseMean > 50) from RNA-seq dataset of 96 hpf EpiCs sorted from *tcf21*^+/+^ and *tcf21*^-/-^ larval hearts.**

